# LDHA maintains the growth and migration of vascular smooth muscle cells and promotes neointima formation via crosstalk between K5 crotonylation and K76 mono-ubiquitination

**DOI:** 10.1101/2023.02.28.530389

**Authors:** Zhi-Huan Chen, Shan-Hu Cao, Zhi-Yan Ren, Han-Mei Jiang, Zhao-Kun Hu, Li-Hua Dong

## Abstract

Phenotypic plasticity of vascular smooth muscle cells (VSMCs) under stress is believed to be a key factor in neointima formation. Lactate dehydrogenase A (LDHA), a key enzyme for glycolysis, has been demonstrated to promote the proliferation and migration of VSMCs. However, the mechanism by which LDHA regulates this process is still unclear. Here we show that the crotonylation and mono-ubiquitination of LDHA are increased in platelet-derived growth factor (PDGF)-BB-induced proliferative VSMCs. Crotonylation at lysine 5 (K5) activates LDHA through tetramer formation to enhance lactate production and VSMCs growth. Mono-ubiquitination at K76 induces the translocation of LDHA into mitochondria, which promotes mitochondria fission and subsequent the formation of lamellipodia and podosomes, thereby enhancing VSMC migration and growth. Furthermore, the increase of crotonylation and ubiquitination were also observed in the carotid arteries of ligation injury mice. Deletion of LDHA K5 crotonylation or K76 mono-ubiquitination decreases ligation-induced neointima formation. Our study reveals a novel mechanism that combines VSMC metabolic reprogramming and behavioral abnormity through crosstalk between LDHA K5 crotonylation and K76 mono-ubiquitination.

## Introduction

As the main pathological process underlying cardiovascular diseases (CVDs), atherosclerosis is the leading cause of mortality among humans throughout the world.^1^ Vascular smooth muscle cells (VSMCs) are the predominant cell type found in medial layer of arteries. Phenotypic change of VSMCs, from a contractile phenotype to a proliferative phenotype, contributes to the development of atherosclerosis and restenosis.^2,3^ Emerging evidence has shown that the phenotypic change of VSMCs is due to a metabolic switch to fulfill the bio-energetic and biosynthetic requirements.^4^ Clinically, atherosclerosis and restenosis have been found to be more common in diabetic patients, suggesting a potential link between inherent abnormalities in glucose metabolism and progression of vascular diseases.^5^ Our studies have previously demonstrated that glucose uptake mediated by glucose transporters 4 (GLUT4) was increased and glucose-6-phosphate dehydrogenase (G6PD) membrane translocation and activation were promoted in PDGF-BB-induced proliferative VSMCs.^6,7^ Other studies have shown that expression of key glycolytic enzymes, Hexokinase 2 (HK2) and lactate dehydrogenase A (LDHA), were increased in PDGF-BB-induced VSMCs and LDHA was reported to play an important role in the proliferation and migration of VSMCs.^8,9^ All the above studies showed that the expression and activity of key enzymes in glucose metabolism possess important roles in the phenotypic change of VSMCs. However, the specific mechanism between glucose metabolism reprogramming and phenotypic change of VSMCs remains unclear.

Protein posttranslational modifications (PTMs) exert a diverse array of biological functions in response to various cellular and extracellular stimuli by modulating enzymatic activity, protein stability, interacting platform, and so on. Lysine crotonylation (Kcr) is a newly discovered PTM originally identified in histone by Tan et al in 2011.^10^ Afterwards, the landscape of these modifications is rapidly expanding and several proteomics studies showed that lysine crotonylation on non-histone proteins also widely exist.^11-13^ Recently, the roles of histone crotonylation in pathophysiologic processes, including kidney disease,^14^ spermatogebesis,^15^ depression,^16^ HIV latency,^17^ cancer^18^ and cardiac homeostasis,^19^ were investigated. Nevertheless, the role of non-histone lysine crotonylation is poorly understood. In our recent study, the interplay of non-histone lysine crotonylome and ubiquitylome in VSMCs phenotypic remodeling were analyzed through bioinformatics.^20^ Given the potential link between glucose metabolism reprogramming and phenotypic switching of VSMCs, here we focused on the key enzymes of glucose metabolism. We find that crotonylation and ubiquitination of LDHA were significantly increased in PDGF-BB-induced proliferative VSMCs.

The Warburg effect or aerobic glycolysis describes an enhanced glycolytic flux and suppressed mitochondrial oxidative phosphorylation even under aerobic conditions in a variety of tumors.^21^ Several recent studies suggested that the Warburg effect is also observed in PDGF-BB-induced proliferative VSMCs.^22,23^ LDHA is a key enzyme of aerobic glycolysis and catalyzes the conversion of pyruvate to lactate with the regeneration of NAD^+^. PTMs of key enzymes in glucose metabolism were shown to participate in the coordination of glycolysis and mitochondrial metabolism. Succinylation mediated mitochondrial translocation of pyruvate kinase M2 (PKM2), which stabilizes VDAC3 and furthermore promotes cell survival under stress.^24^ O-GlcNAcylation induces PGK1 translocation into mitochondria to inhibit pyruvate dehydrogenase (PDH) complex to reduce oxidative phosphorylation.^25^ Mitofusin 2 (MFN2) prevents neointimal hyperplasia in vein grafts via decreasing phosphofructokinase 1 (PFK1) ubiquitination, which increases glycolysis and decreases oxidative phosphorylation.^26^ Previous observation that targeting LDHA by shRNA stimulated oxidative phosphorylation of cancer cells provides a direct link between glycolysis and oxidative phosphorylation that involves LDHA.^27^ Phosphorylation, acetylation and succinylation of LDHA were also demonstrated to be involved in the regulation of aerobic glycolysis in different type of tumors.^28-30^ These findings prompted us to investigate whether crotonylation and ubiquitination of LDHA observed in our previous study play important roles in the regulation of glucose metabolism and the phenotypic change of proliferative VSMCs.

In this study, we found that ‘crosstalk’ between ubiquitination and crotonylation of LDHA participated in the phenotypic switching of VSMCs. We showed that LDHA is crotonylated at lysine 5 (K5) and crotonylation of LDHA K5 enhances the formation of active, tetrameric LDHA, support cell growth. Simultaneously, LDHA is mono-ubiquitinated at lysine 76 (K76) and mono-ubiquitination of LDHA K76 promoted mitochondrial division, VSMC migration and cell growth. Our research illustrated a new mechanism underlying aerobic glycolysis in proliferative VSMCs and provided a new target for the clinical treatment of CVDs.

## Results

### LDHA expression is important for VSMC growth and migration

In our recent study, we performed the modified omics and proteomic analysis of VSMC phenotypic switching stimulated with PDGF-BB. The crotonylated and ubiquitinated pan-antibody was used to enrich proteins and then subjected to a high-throughput mass spectrometry analysis. We found that enzymes of the glucose metabolism pathways including glycolysis/gluconeogenesis, pentose phosphate pathway, pyruvate metabolism and TCA cycle, were modified extensively by crotonyl and ubiquitin.^20^ The modification of the enzymes might modulate the pathways directly or indirectly. Among these enzymes, both the crotonylation and ubiquitination of LDHA were significantly increased in PDGF-BB-induced VSMCs, following that the expression of LDHA was also upregulated, thus we screen LDHA as an important enzyme that may regulate VSMC phenotypic switching.

To verify this finding, cultured VSMCs *in vitro* were stimulated with PDGF-BB (10 ng/mL) for different durations (0-48 h). The results showed that the expression of LDHA was significantly increased at 12 h and peaked at 24 h on PDGF-BB stimulation, which paralleled the VSMC proliferative/synthetic state, as evidenced by alterations in the expression of cell proliferative marker (PCNA) and VSMC contractile marker (SM22α)^31-33^ (Figure 1A). At the *in vivo* level, LDHA expression was gradually induced in injured carotid arteries compared with sham arteries, as indicated by assessment at days 7, 14 and 21 after ligation injury/sham procedure; in parallel, reduction in SM22α and increasement in PCNA were also observed (Figure 1B). LDHA catalyzes the reduction of pyruvate to lactate, we further showed that the lactate generation paralleled the change of increased LDHA expression in PDGF-BB-induced VSMCs (Figure 1C), which indicated the LDHA activity and the glycolysis pathway were induced in proliferative VSMCs.

**Figure 1.**
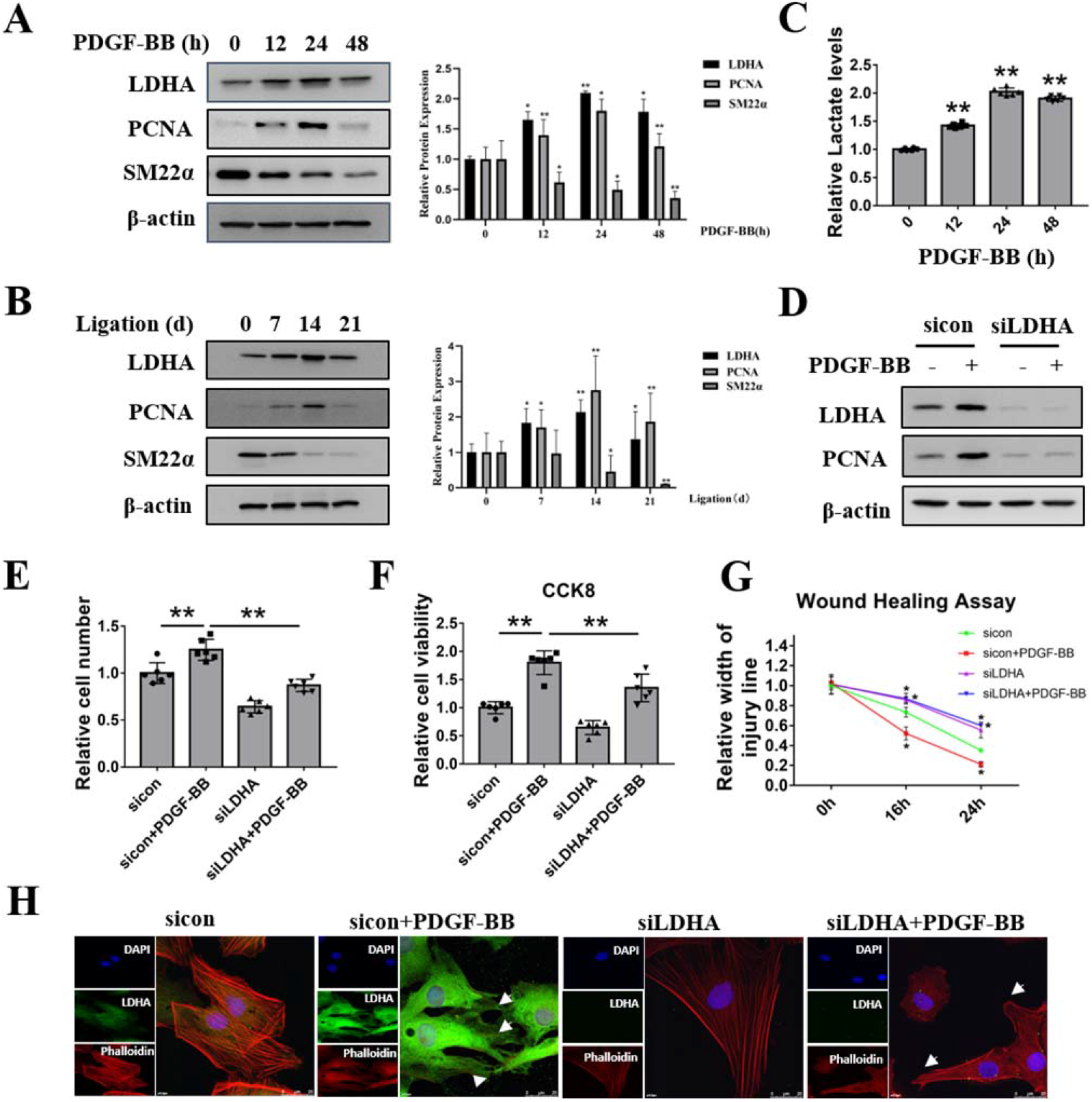
LDHA expression is important for VSMC growth and migration. (A) Immunoblotting analysis of LDHA protein expression in VSMCs stimulated by PDGF-BB for different durations (12, 24 and 48 h). (B) Immunoblotting analysis of LDHA protein expression in injured carotid arteries at days 7, 14 and 21 after ligation injury/sham procedure. (C) Quantitative analysis of lactate levels in VSMCs stimulated by PDGF-BB for different durations. (D) Immunoblotting analysis of PCNA protein expression upon depletion of LDHA using sicon/siLDHA with (+) or without (-) PDGF-BB stimulation (24 h). (E) Cell growth of VSMCs upon depletion of LDHA using sicon/siLDHA with (+) or without (-) PDGF-BB stimulation (24 h). (F) Cell viability of VSMCs upon depletion of LDHA using sicon/siLDHA with (+) or without (-) PDGF-BB stimulation (24 h). (G) Wound-healing assay in VSMCs upon depletion of LDHA using sicon/siLDHA with (+) or without (-) PDGF-BB stimulation (24 h). Quantitation of relative width of injury line was shown. (H) Immunofluorescence staining analysis of LDHA and actin filaments with phalloidine (pha) in VSMCs upon depletion of LDHA using sicon/siLDHA with (+) or without (-) PDGF-BB stimulation (24 h). The scale bar was 25 μm.

To determine the possible role of LDHA in phenotypic switching of VSMCs, we depleted endogenous LDHA in cultured VSMCs with or without PDGF-BB treatment by a specific small interfering RNA (siRNA). PCNA expression was no longer increased with the treatment of PDGF-BB after depletion of LDHA (Figure 1D). Moreover, VSMC numbers were increased after PDGF-BB treatment but deficiency of LDHA repressed the PDGF-BB induced growth advantage (Figure 1E). CCK8 analysis showed that cell viability was significantly increased after PDGF-BB stimulation in VSMCs while this elevation was reduced by LDHA silencing (Figure 1F). These results highlight the importance of LDHA in VSMC growth from PDGF-BB stimulation.

VSMC migration, a well-known characteristic of phenotype switching, is an early event during the formation of atherosclerotic lesions.^34^ Wound-healing assay showed that knockdown of LDHA decreased PDGF-BB-induced VSMC migration (Figure 1G and Supplementary material online, Figure S1). Migration generally begins with the extension of protrusions at the leading edge of the cells and this membrane protrusion is called lamellipodia.^35^ Therefore, we observed the effect of LDHA in lamellipodia formation using phalloidin, a “gold standard” for actin filament staining. The results showed lamellipodia formation was elevated after PDGF-BB stimulation in VSMCs. While this effect was not observed in LDHA-silenced VSMCs with or without PDGF-BB treatment (Figure 1H), indicating that LDHA is perhaps involved in the effect of PDGF-BB on lamellipodia formation. Moreover, we also observed podosome, a structure that is crucial for migration, in PDGF-BB-induced VSMCs but not in the control group or LDHA-silenced groups (Figure 1H). These data suggest that LDHA is essential to the podosome formation, which related with the lamellipodia and VSMC migration and that LDHA is functionally important in proliferative VSMCs.

### LDHA is highly crotonylated at lysine 5 in proliferative/synthetic VSMCs

Following the initial discovery of lysine crotonylation, histone crotonylation has become intensively studied over the last several years.^14-19^ However, the functional impact of non-histone Kcr is still in the infancy stage. To confirm the crotonylome results, immunoprecipitation was performed utilizing a pan anti-crotonyl-lysine antibody at the *in vitro* and *in vivo* level. As expected, crotonylation level of LDHA was increased at 12 h and peaked at 24 h on PDGF-BB stimulation, which is consistent with the VSMC proliferative/synthetic state (Figure 2A). In injured carotid arteries, LDHA crotonylation also paralleled the intimal hyperplasia degree which was assessed by ligation for 7, 14 and 21 days (Figure 2B). These results suggested that LDHA crotonylation is perhaps involved in the VSMC phenotypic switching.

**Figure 2.**
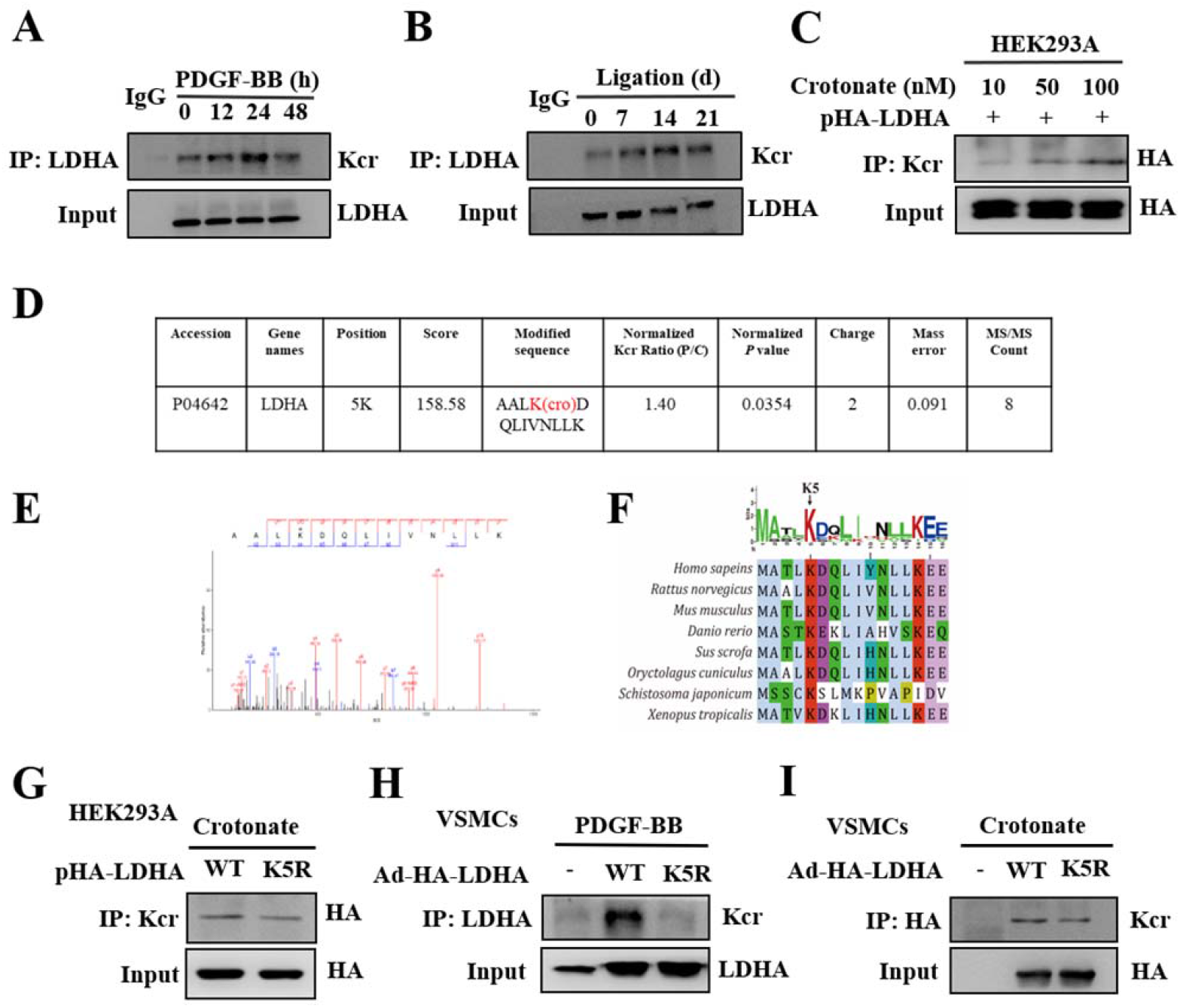
LDHA is highly crotonylated at lysine 5 in proliferative/synthetic VSMCs. (A) Complexes immunoprecipitated (IP) with anti-crotonyl-lysine antibody were immunoblotted (IB) with anti-LDHA antibodies in VSMCs with (+) PDGF-BB treatment for different durations (12, 24 and 48 h). (B) Analysis of LDHA crotonylation in injured carotid arteries at days 7, 14 and 21 after ligation injury/sham procedure. (C) Analysis of LDHA crotonylation in HEK293A cells transfected with HA-LDHA constructs followed by different concentrations of crotonate treatment. (D and E) Identification of crotonylated LDHA peptides by mass spectrometry analysis of VSMCs upon PDGF-BB stimulation. VSMCs without PDGF-BB treatment were examined as the control group. (F) Sequence analysis of LDHA K5 from *Xenopus tropicalis* to *Homo sapeins* using the software of Bioedit. (G) Analysis of LDHA crotonylation in HEK293A cells transfected with HA-LDHA or HA-K5R constructs followed by crotonate treatment. (H) Analysis of LDHA crotonylation in VSMCs transduced with the LDHA WT or K5R adenovirus followed by PDGF-BB stimulation. (I) Analysis of LDHA crotonylation in VSMCs transduced with the LDHA WT or K5R adenovirus followed by crotonate treatment.

Increasing evidences have shown that adding crotonate to cell cultures leads to an increase in lysine crotonylation through the direct production of crotonyl-CoA.^36^ Here we synthesized and constructed the LDHA full-length plasmid (HA-LDHA) and HEK293 cells were transfected with HA-LDHA constructs followed by different concentrations of crotonate stimulation. Complexes immunoprecipitated with pan anti-Kcr antibodies were immunoblotted with HA antibodies. Consistently, the result showed a dose-dependent increase of LDHA crotonylation (Figure 2C).

In our crotonylome, mass spectrometry analysis showed that LDHA was crotonylated on lysine 5 (K5) in PDGF-BB-induced VSMCs, where the level of LDHA crotonylation was 1.4 times higher than in quiescent cell (Figure 2D and E). The sequence analysis showed that the K5 residue is well conserved from *schistosoma japonicum* to mammals (Figure 2F). In order to ensure the crotonylation site, we mutated K5 of LDHA full-length plasmid (HA-LDHA) to arginine (R) HA-LDHA-K5R (mimicking deletion). HEK293 cells were transfected with HA-LDHA and HA-K5R constructs followed by crotonate treatment. K5R mutant displayed a significant decrease in LDHA crotonylation compared to the WT, indicating that K5 is the site for crotonylation of LDHA (Figure 2G). To further confirm our conclusion, we transduced the LDHA K5 site-mutant adenovirus Ad-HA-LDHA K5R into VSMCs followed by PDGF-BB or crotonate stimulation. Immunoblotting with the anti-LDHA antibody showed the similar results (Figure 2H and I). We speculated that K5 might be a key and evolutionarily preserved site for regulating LDHA activity in a crotonyl-dependent manner.

### LDHA K5 crotonylation promotes tetramer formation and VSMC growth

LDHA is a globular protein of mixed α/β secondary structure with a protruding N-terminal segment which was shown to possess an important role in tetramerization to establish active form of the enzyme.^37,38^ Docking analysis of LDHA using PyMol software showed that within each tetramer, K5 is highly exposed and extends outside and the extended N-terminus of each monomer makes pairwise interactions with two adjacent molecules (Figure 3A). Crotonylation at this site could therefore affect the binding capacity and protein-protein interactions, thereby influencing the structure and function of the protein.

**Figure 3.**
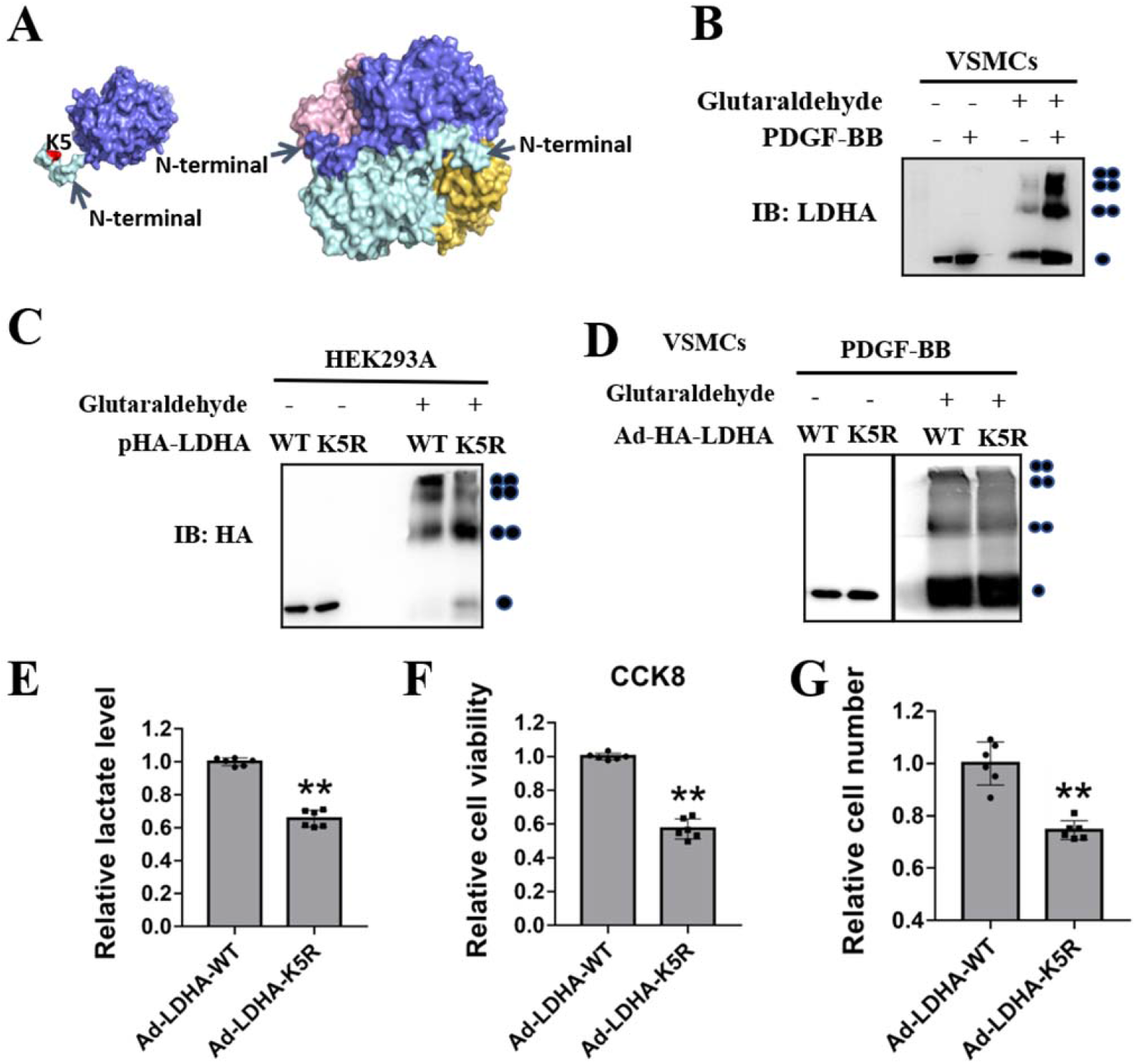
LDHA K5 crotonylation promotes tetramer formation and VSMC growth. (A) Docking analysis of LDHA using PyMol software. The sequence was obtained from PBD (DOI: 10.2210/pdb4AJJ/pdb). (B) Analysis of LDHA tetramerization in cells using a glutaraldehyde crosslinked method. VSMCs with (+) or without (-) PDGF-BB stimulation were crosslinked by 0.05% glutaraldehyde and analyzed by western blotting using LDHA antibody. Tetrameric, dimeric, and monomeric LDHA were indicated. (C) Analysis of LDHA tetramerization in HEK293A cells transfected with HA-LDHA or HA-K5R constructs. (D) Analysis of LDHA tetramerization in VSMCs transduced with the LDHA WT or K5R adenovirus followed by PDGF-BB stimulation. (E) Quantitative analysis of lactate levels in VSMCs transduced with the LDHA WT or K5R adenovirus followed by PDGF-BB stimulation. (F) Cell viability of VSMCs transduced with the LDHA WT or K5R adenovirus followed by PDGF-BB stimulation. (G) Cell growth of VSMCs transduced with the LDHA WT or K5R adenovirus followed by PDGF-BB stimulation.

Protein crosslinking assay is usually used to identify enzyme oligomerization, dimerization and tetramerization.^39,40^ Here VSMCs with (+) or without (-) PDGF-BB treatment were crosslinked by 0.05% glutaraldehyde and analyzed by western blotting using LDHA antibody. The result showed that the formation of active, tetrameric LDHA was significantly increased in PDGF-BB induced proliferative VSMCs, positively correlated with the induction of crotonylation levels (Figure 3B). Since crotonate treatment promotes LDHA crotonylation in VSMCs, to further verify the relationship between crotonylation levels and tetramerization of LDHA, we added different concentrations of crotonate to VSMC cultures and examined LDHA tetramerization. Consistently, the formation of active, tetrameric LDHA increased in a similar manner to that of crotonylation levels (Supplementary material online, Figure S2A).

To further corroborate the result that crotonylation at K5 could affect the active form of LDHA, we determined the tetramerization of LDHA WT and K5R mutant in HEK293 cells. Overexpression of K5R mutant displayed a significant decrease in LDHA tetramerization, indicating that crotonylation at K5 may mediate the formation of active, tetrameric LDHA (Figure 3C). To further verify this point, VSMCs were infected with the LDHA K5 site-mutant adenovirus Ad-HA-K5R followed by PDGF-BB or crotonate stimulation. The results were consistent with the above (Figure 3D and Supplementary material online, Figure S2B). Collectively, these findings suggest that crotonylation at K5 enhances the formation of active, tetrameric LDHA.

To explore the impact of LDHA crotonylation on VSMC growth, we examined the level of lactate, a direct product of LDHA, in VSMCs which were transduced with Ad-HA-LDHA WT or K5R followed by PDGF-BB stimulation. The results showed that lactate production was downregulated in LDHA K5R expressing VSMCs compared to the LDHA WT expressing cells upon PDGF-BB stimulation (Figure 3E). Correspondingly, in keeping with the lactate level, cell viability, cell numbers and PCNA expression exhibited a similar decrease in LDHA K5R expressing cells compared with the LDHA WT expressing cells upon PDGF-BB stimulation (Figure 3F and G, Supplementary material online, Figure S2C). Thus, our data indicate that LDHA K5 crotonylation is functionally important for VSMC growth.

### LDHA is mono-ubiquitinated at K76

To confirm the ubiquitylome results, immunoprecipitation with anti-Ub antibody was performed in PDGF-BB induced VSMCs. A mono-ubiquitinated form of LDHA was detected and ubiquitination level of LDHA was upregulated at 12 h and peaked at 24 h on PDGF-BB stimulation, which is also consistent with the VSMC proliferative/synthetic state (Figure 4A). At the *in vivo* level, LDHA ubiquitination paralleled the intimal hyperplasia degree which was assessed by ligation for 7, 14 and 21 days, suggesting that LDHA mono-ubiquitination may play a key role in VSMC phenotypic switching (Figure 4B). In our ubiquitylome analysis, ubiquitination of LDHA in PDGF-BB induced proliferative VSMCs was predicted more than 1.3 times higher than in the control group and two putative ubiquitination sites of LDHA, K5 and K76, were identified by mass spectrometry (Figure 4C and D). Sequence analysis showed that both K5 and K76 are evolutionarily conserved from *schistosoma japonicum* to mammals (Figure 2E and 4E). To clarify the major ubiquitination site of LDHA, HEK293 cells were transfected with His-Ub along with HA-LDHA-WT, HA-LDHA-K5R or HA-LDHA-K76R plasmids. Substitution of K76, but not K5, dramatically decreased the LDHA mono-ubiquitination to the greatest degree (Figure 4F), indicating that K76, is a major mono-ubiquitination site in LDHA. To further verify our conclusion, VSMCs were transduced with Ad-HA-LDHA WT, K5R or K76R followed by PDGF-BB stimulation. The results showed that overexpression of the LDHA K76R mutant, but not K5R, exhibited a remarkable decrease of LDHA ubiquitination (Figure 4G). Taken together, these results suggest that LDHA is mono-ubiquitinated at K76 in proliferated VSMCs and may possess an important role in VSMC phenotypic switching.

**Figure 4.**
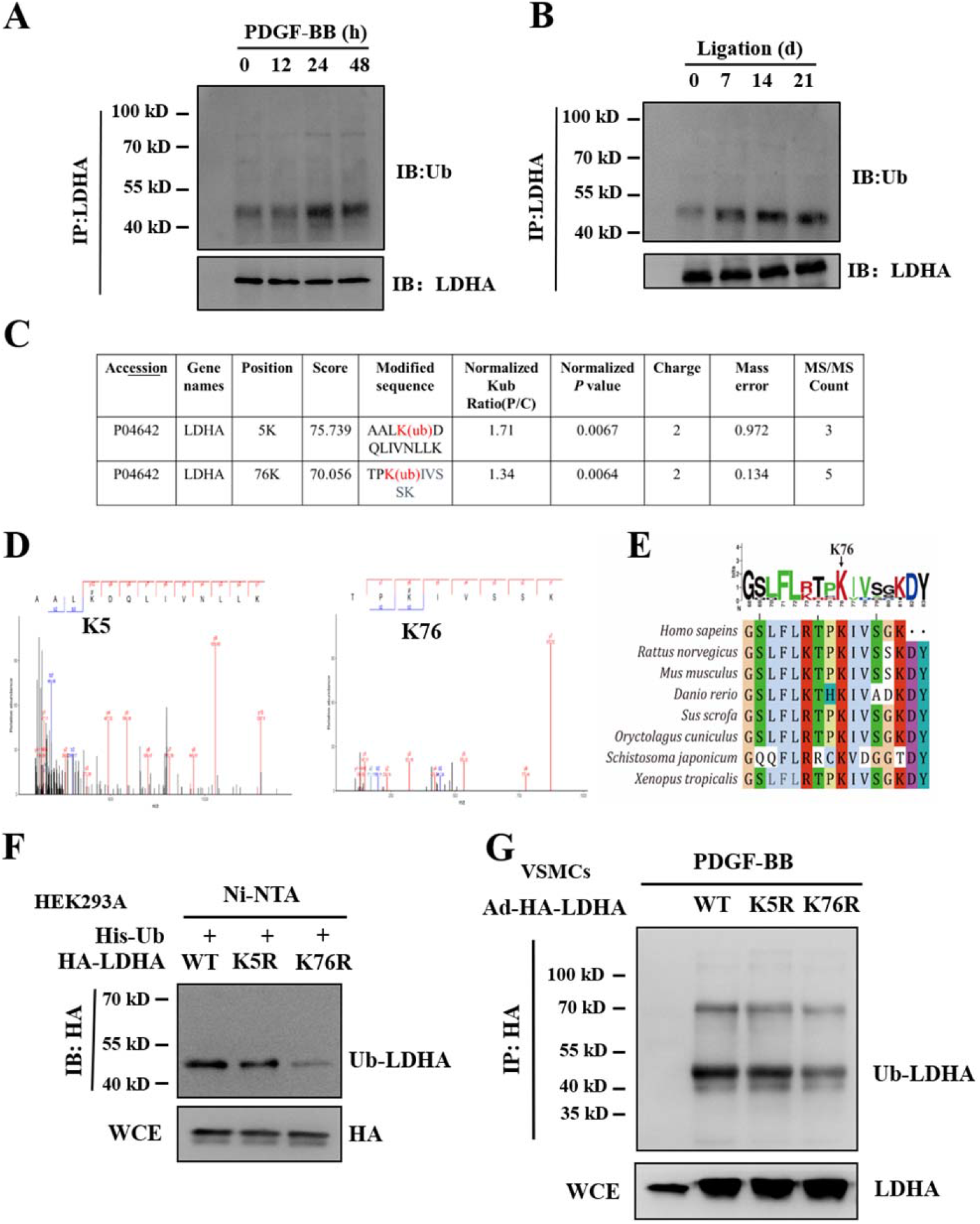
LDHA is mono-ubiquitinated at K76. (A) Complexes immunoprecipitated (IP) with anti-LDHA antibodies were immunoblotted (IB) with ubiquitin antibodies in VSMCs with (+) PDGF-BB treatment for different durations (12, 24 and 48 h). (B) Analysis of LDHA ubiquitination in injured carotid arteries at days 7, 14 and 21 after ligation injury/sham procedure. (C and D) Identification of ubiquitinated LDHA peptides by mass spectrometry analysis of VSMCs upon PDGF-BB stimulation. VSMCs without PDGF-BB treatment were examined as the control group. (E) Sequence analysis of LDHA K76 from *Xenopus tropicalis* to *Homo sapeins* using the software of Bioedit. (F) Analysis of LDHA ubiquitination in HEK293A cells transfected with His-Ub along with HA-LDHA-WT, HA-LDHA-K5R or HA-LDHA-K76R plasmids using Ni-NTA agarose beads. (G) Analysis of LDHA ubiquitination in VSMCs transduced with the LDHA WT, K5R or K76R adenovirus followed by PDGF-BB stimulation.

### Mono-ubiquitination of LDHA K76 promotes mitochondrial division, VSMC migration and growth

Since PTMs of major glycolytic enzymes were reported to play pivotal roles in coordinating glycolysis and mitochondrial metabolism,^24-26^ we next set up to investigate whether PTMs of LDHA is involved in this effect. We first evaluated the relationship between LDHA and mitochondrial through immunofluorescence staining. VSMCs were co-stained with MitoTracker red, a fluorescent marker for mitochondria, and anti-LDHA antibody. The results showed that LDHA co-localized with mitochondria after PDGF-BB stimulation (Figure 5A). Mitochondria are morphologically dynamic, undergoing continuous shape change through fission and fusion.^41^ Here we observed that mitochondria were scattered in the cytoplasm in a fused state in the quiescent VSMCs while were divided, shortened and became smaller in the proliferative VSMCs induced by PDGF-BB (Figure 5A). To further confirm the translocation of LDHA into mitochondria, cell fractionation and immunoblotting were analyzed. Expression of LDHA was greatly elevated in the mitochondrial fraction of VSMCs treated with PDGF-BB (Figure 5B). To address whether LDHA modulates the fission and fusion of mitochondria, we silenced LDHA by siRNA in VSMCs. Knockdown of LDHA abolished the PDGF-BB induced mitochondria fission, suggesting that LDHA play a key role in the regulation of mitochondria morphology (Figure 5C).

**Figure 5.**
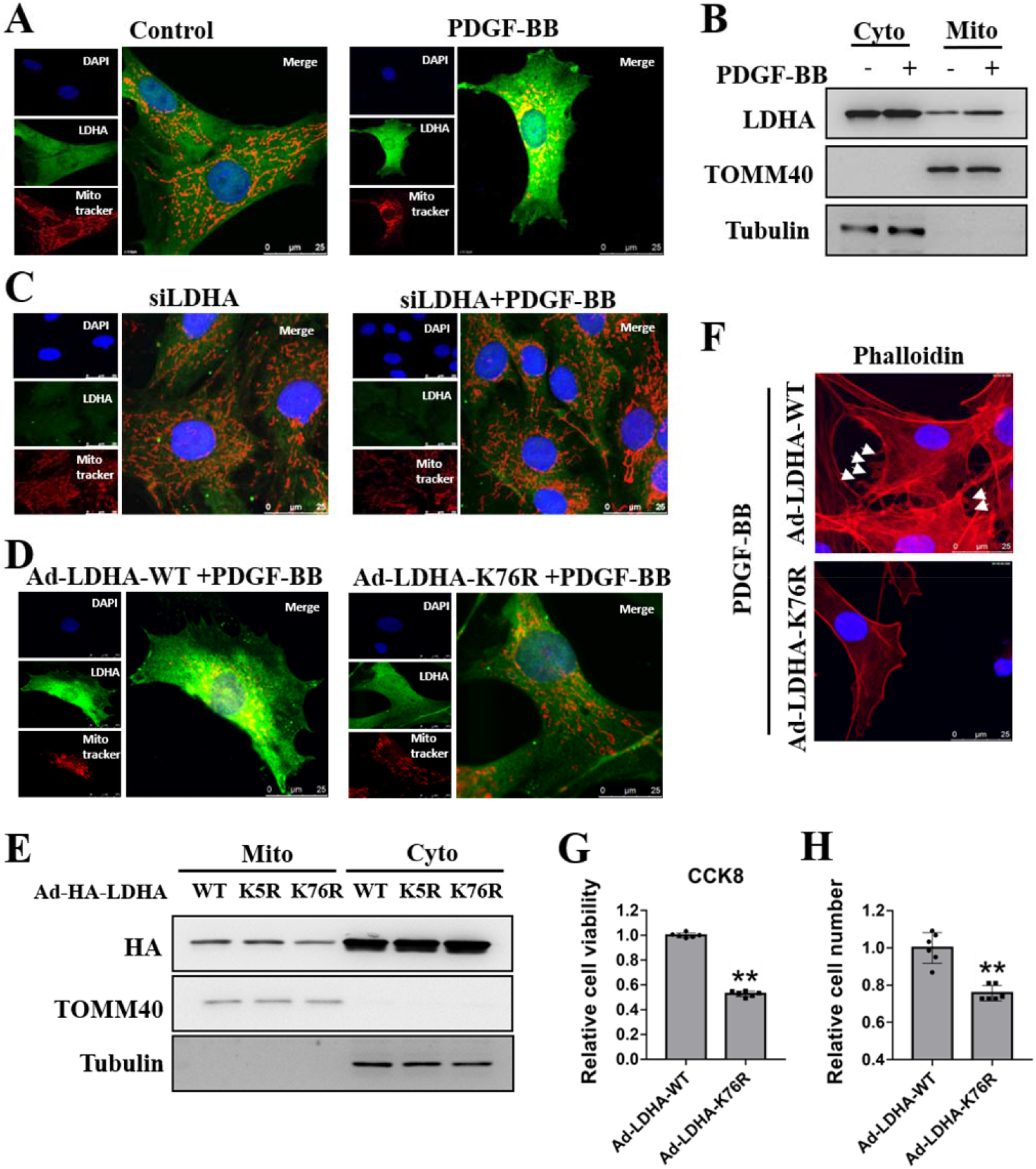
Mono-ubiquitination of LDHA K76 promotes mitochondrial division, VSMCs migration and growth. (A) Immunofluorescence staining analysis of LDHA and mitochondria with Mito tracker in VSMCs with (+) or without (-) PDGF-BB stimulation. The scale bar was 25 μm. (B) LDHA expression level in the mitochondria fractions which were extracted from VSMCs with (+) or without (-) PDGF-BB treatment. (C) Immunofluorescence staining analysis of LDHA and mitochondria in VSMCs upon depletion of LDHA using siLDHA with (+) or without (-) PDGF-BB stimulation. (D) Immunofluorescence staining analysis of LDHA and mitochondria in VSMCs transduced with the LDHA WT or K76R adenovirus followed by PDGF-BB stimulation. (E) LDHA expression level in the mitochondria fractions which were extracted from VSMCs transduced with the LDHA WT, K5R or K76R adenovirus followed by PDGF-BB stimulation. (F) Immunofluorescence staining analysis of actin filaments with phalloidine (pha) in VSMCs transduced with the LDHA WT or K76R adenovirus followed by PDGF-BB stimulation. The scale bar was 25 μm. (G) Cell viability of VSMCs transduced with the LDHA WT or K76R adenovirus followed by PDGF-BB stimulation. (H) Cell growth of VSMCs transduced with the LDHA WT or K76R adenovirus followed by PDGF-BB stimulation.

Ubiquitination is well established as a canonical signal for degradation by the ubiquitin-proteasome system (UPS).^42^ Intriguingly, here PDGF-BB induced elevation of ubiquitination did not decrease the protein level of LDHA in VSMCs, suggesting that mono-ubiquitination of LDHA serves a non-degradative function. Emerging evidence has shown that mono-ubiquitination is related to transcription,^43^ localization^44^ or function of the protein.^45^ Considering the co-localization of LDHA with mitochondria after PDGF-BB treatment in VSMCs, we assume that mono-ubiquitination perhaps mediated the translocation of LDHA into mitochondria. To confirm our hypothesis, we transduced Ad-HA-LDHA WT and K76R into VSMCs followed by PDGF-BB stimulation to track the cellular distribution of LDHA. Overexpression of K76R LDHA resulted in reduced LDHA translocation to mitochondria, accompanied by decreased mitochondrial fission (Figure 5D). The subsequent mitochondrial fractionation and immunoblotting analysis further confirmed the immunostaining results (Figure 5E). These findings suggested that K76 mono-ubiquitination mediates mitochondrial translocation of LDHA and thus promotes mitochondrial fission.

A previous study has demonstrated that mitochondrial fission contributes to VSMC migration.^46^ To gain a better understanding of the impact of LDHA mono-ubiquitination on VSMC migration, we assessed the lamellipodia formation by phalloidin staining in VSMCs which were transduced with Ad-HA-LDHA WT or K76R followed by PDGF-BB stimulation. The results showed that LDHA WT expressing cells exhibited an enhanced ability of lamellipodia formation after PDGF-BB treatment. Whereas in LDHA K76R expressing cells, lamellipodia were not detected (Figure 5F). Consistently, we also observed podosomes, another confirmation for migration, in LDHA WT expressing cells but not in LDHA K76R expressing cells (Figure 5F). In addition, VSMC viability and numbers were increased in LDHA WT expressing cells while were decreased in LDHA K76R expressing cells (Figure 5G and H). Collectively, these findings indicate that mono-ubiquitination of LDHA K76 is critical for VSMC migration and growth.

### LDHA crotonylation and ubiquitination exacerbates post-injury neointima formation *in vivo*

To investigate whether LDHA crotonylation and ubiquitination contributes to intimal hyperplasia *in vivo*, we performed periadventitial infection of adenoviruses encoding LDHA WT, K5R or K76R in carotid arteries. Fourteen days after ligation, adenovirus-mediated overexpression of LDHA WT, K5R or K76R were verified. Consistently, LDHA overexpression in carotid arteries significantly exacerbated ligation-induced neointima formation (Figure 6A). The neointima area and ratio of neointima to media area was significantly greater in Ad-LDHA-infected arteries than that in WT arteries (Figure 6B and C). However, there was no difference in the media area or circumference of the external elastic lamina (Figure 6D and E). In accordance, LDHA overexpression promoted the induction of VSMC proliferative marker PCNA expression after ligation injury *in vivo* (Supplementary material online, Figure S3). In contrast, the intimal hyperplasia degree is obviously decreased in LDHA K5R and K76R expressing arteries compared to the LDHA WT expressing arteries (Figure 6A-C). In parallel, reduction in PCNA was also observed in LDHA K5R and K76R expressing arteries (Supplementary material online, Figure S3). These results showed that LDHA K5 crotonylation and K76 mono-ubiquination promote intimal hyperplasia of injuried arteries.

**Figure 6.**
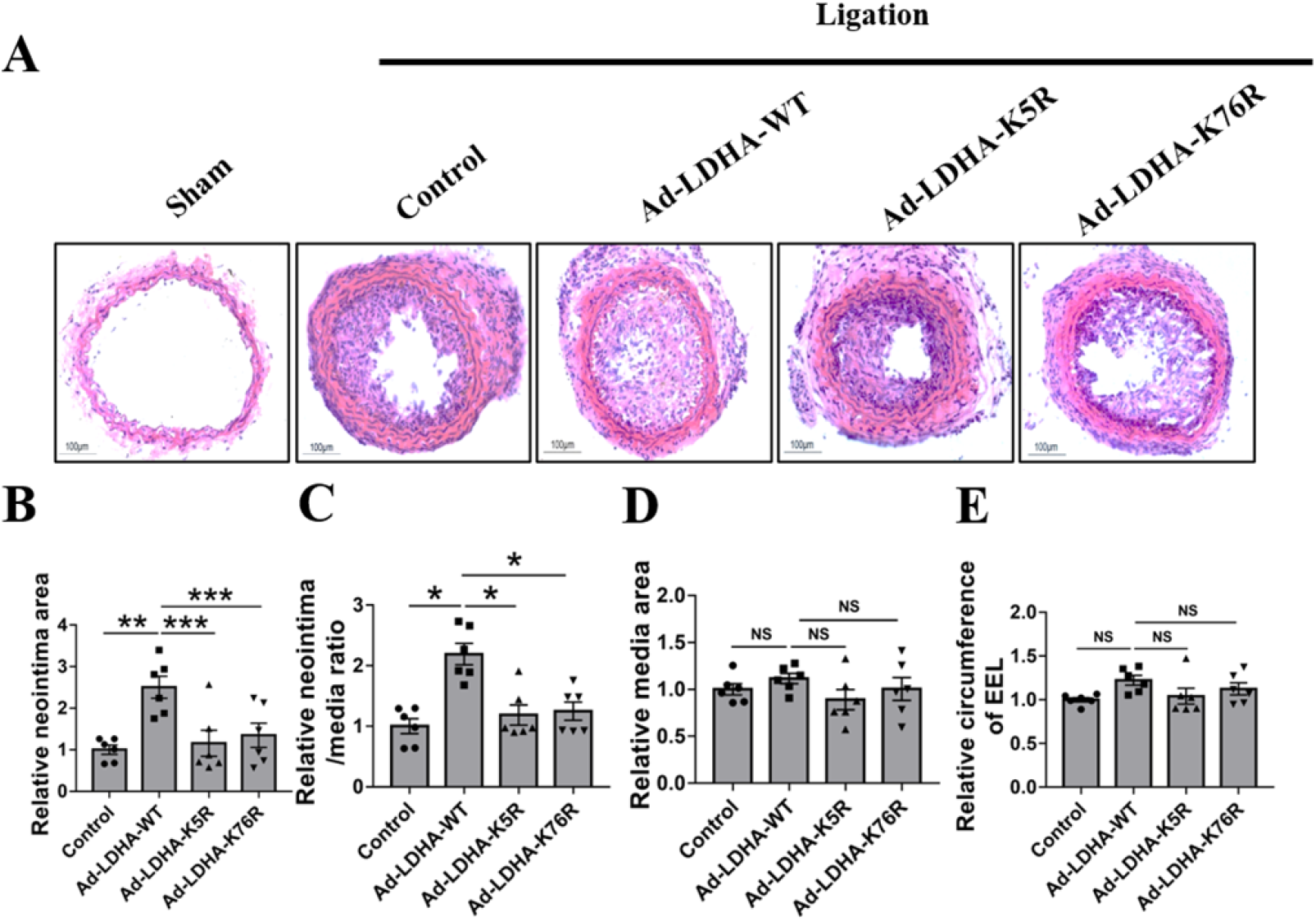
Mono-ubiquitination of LDHA K76 promotes mitochondrial division, VSMCs migration and growth. (A) H&E staining analysis of wire-injured carotid arteries transduced with the LDHA control, WT, K5R or K76R adenovirus at day 14. Quantitative analysis of (B) the intima area, (C) intima-to-media ratio, (D) media area, and (E) external elastic lamina (EEL) circumference in wire-injured carotid arteries transduced with the LDHA control, WT, K5R or K76R adenovirus at day 14.

## Discussion

Metabolic reprogramming characterized by an increased aerobic glycolysis and suppressed oxidative phosphorylation is a typical feature of proliferative cells. LDHA, a key metabolic enzyme in glycolysis, is upregulated in many proliferative cells and promotes cell survival through multiple pathways, including LDHA expression,^47^ posttranslational modification^28-30^, localization^39^ and protein interactions.^48^ Recently, it has been shown that LDHA is indispensable for VSMC proliferation and migration to maintain the atherogenic phenotype.^9^ However, mechanisms underlying such a role are still unknow. Here, we uncover a novel biochemical mechanism that involves both glycolytic and nonglycolytic functions of LDHA in regulating VSMC metabolic reprogramming and phenotypic switching. Crotonylation at K5 activates LDHA through enhancing the formation of tetramer, which contributes to the production of lactate and subsequent promotes VSMC growth under PDGF-BB stimulation. Mono-ubiquitination at K76 regulates the translocation of LDHA into mitochondria, which promotes mitochondria fission and subsequent the formation of lamellipodia and podosomes, thereby enhancing VSMC migration and growth under PDGF-BB stimulation (Figure 7).

**Figure 7.**
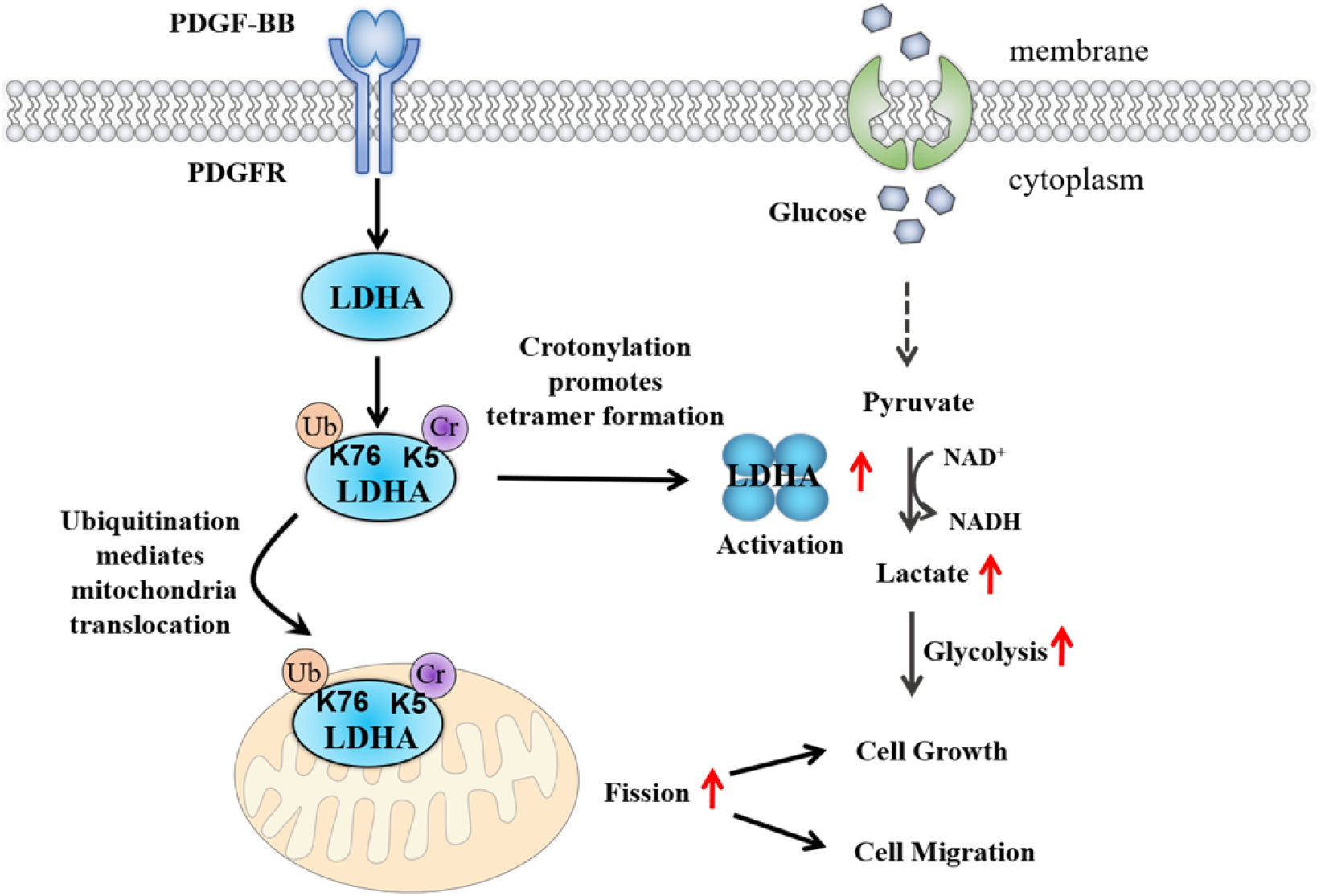
Crosstalk between LDHA K5 crotonylation and K76 mono-ubiquitination in regulating VSMC phenotypic switching. Crotonylation at K5 activates LDHA enzymatic activity and mono-ubiquitination at K76 mediates the translocation of LDHA into mitochondria. Both glycolytic and nonglycolytic functions of LDHA maintain the growth and migration of VSMCs and promote neointima formation.

Crotonylation of proteins is a recently discovered PTM that occurs leadingly on the lysine residue. A growing body of study has implied that short-chain fatty acids including crotonate and their cognate acylations participate in CVDs.^49^ In this study, we found that lysine crotonylation is widespread on proteins of diverse functions and localizations in PDGF-BB induced proliferative VSMCs. Glucose metabolism is vital for cell survival and function because it provides substrates and energy for numerous enzymatic reactions. Our study shows that LDHA is elevated in PDGF-BB induced proliferative VSMCs. Silencing of LDHA in cultured VSMCs suppresses cell growth from PDGF-BB stimulation *in vitro*, while overexpression of LDHA is sufficient to drive VSMC growth and intimal hyperplasia. Furthermore, we find that LDHA crotonylation is significantly increased in PDGF-BB induced proliferative VSMCs. Mechanistically, crotonylation dependent tetramer formation of LDHA enhances the activity of LDHA and the formation of lactate, promoting the rapid growth of VSMCs. Our study reveals an important mechanism by which LDHA regulates VSMC phenotypic switching through crotonylation.

The three-dimensional structure of LDHA indicates that lysine 5 is located in the N-terminal alpha-helix region of LDHA.^29^ N-terminal arms of each subunit have an important role in enzyme tetramerization to establish active form of the enzyme.^37^ In this study, we identified K5 as an important regulatory crotonylation site within the LDHA protein, as PDGF-BB–induced tetramer formation of LDHA was abolished by overexpression of the LDHA K5R mutant. Moreover, the PDGF-BB–induced lactate formation and VSMC growth were simultaneously decreased in LDHA K5R expressing cells. Thus, we propose that crotonyl modification at this site enables LDHA to acquire the ability of facilitating LDHA activation.

Protein ubiquitination is an important PTM that regulates various biological functions.^42^ Ubiquitination involves the covalent attachment of the 76-amino acid eukaryotic molecule ubiquitin (Ub) to substrate proteins, which can result in protein mono-ubiquitination or polyubiquitination.^50^ Studies on polyubiquitination mainly focused on the K-48 and K-63 linked chains. K-48 linked polyubiquitination participates in the canonical signal for proteasomal degradation and K-63 linked polyubiquitination plays important roles in cellular processes, including DNA repair, signal transduction, receptor endocytosis and localization.^7,51,52^ Unlike polyubiquitin, mono-ubiquitinated proteins are usually stable and hard to degrade. In general, mono-ubiquitinated target proteins have the functions of regulating transcriptional, gene expression, protein activation and localization. Histone H2AK119 mono-ubiquitination is essential for polycomb-mediated transcriptional repression.^43^ Mono-ubiquitination of RPS27a controls the maturation of the 40S ribosomal subunit.^53^ Proteasomal stress-induced mono-ubiquitination of PINK1 mediates PINK1 nuclear translocation.^54^ In this study, we show that LDHA is mono-ubiquitinated in PDGF-BB induced proliferative VSMCs and we identified K76 as the ubiquitination site. A previous study has shown that LDHA is mono-ubiquitinated in skeletal muscle cells exposed to oxidative stress.^55^ However, the underlying mechanism has not been fully investigated. Here, LDHA is mono-ubiquitinated at K76 in proliferated VSMCs. suggesting that mono-ubiquitinated LDHA mediates a non-degradative function.

Traditionally, mitochondrial metabolism is believed to be the most energetically efficient way of producing ATP in the presence of oxygen. However, a growing number of concepts show that ATP can be produced in large amounts outside of the mitochondria in the cytoplasm, enough to support even rapidly proliferating cancer cells.^56^ Vasculature is a tissue that does not require large amounts of energy. A recent review show that acquired abnormalities of mitochondrial metabolism and dynamics promote a cancer like hyperproliferative phenotype in all pulmonary artery smooth muscle cells (PAMSC).^57^ Mitochondrial dynamics refers to the morphology network comprising individual organelles that continuously join (fusion) and fragment (fission). Li et al showed that decreasing mitochondrial fission diminishes VSMC migration and ameliorates intimal hyperplasia.^46^ Another study reported that mitochondrial fission is a prerequisite for PAMSC proliferation.^58^ These findings indicate that mitochondrial fission facilitates VSMC proliferation and migration. However, a direct correlation between mitochondrial morphology and VSMC phenotypic switching remains ill-defined. In this study, we find that LDHA translocates into mitochondrial by mono-ubiquitin modification after PDGF-BB stimulation in VSMCs, which promotes mitochondrial fission, thereby facilitating VSMC proliferation and migration. In addition to LDHA nuclear and lysomomal translocation, our study is the first time that describes mitochondrial translocation of LDHA.

In summary, we demonstrated that LDHA is activated in response to PDGF-BB stimulation in VSMCs accompanied by the increased translocation of LDHA from the cytosol to mitochondrial. The coordination of glycolytic enzyme activity and mitochondrial dynamics accounts for the hyper-proliferative phenotype in VSMCs and intimal hyperplasia. Our study provides potential therapeutic targets for CVDs in the pathway that is related to LDHA. Studies are ongoing in our lab to investigate metabolic enzymes in proliferative VSMCs to provide more therapeutic targets.

## Materials and Methods

### Cell culture and treatment

VSMCs were isolated from the thoracic aorta of 60-80g male Sprague-Dawley rats. Cells were cultured in low-glucose Dulbecco’s Modified Eagle Medium (DMEM) (Invitrogen, US) supplemented with 10% fetal bovine serum (FBS), 100 U/mL penicillin and 100 µg/mL streptomycin, in a humidified incubator at 37℃ with an atmosphere of 5% CO_2_ and only passages 3 to 5 were used in the experiments. The cells at 70-80% confluence were treated with PDGF-BB (10 ng/mL) or crotonate for further culture. HEK293A cells were cultured in high glucose DMEM with 10% FBS.

### Carotid artery ligation model in mice and adenovirus transduction

The left carotid artery of male C57BL/6J mice was ligated with a 6-0 silk suture before the common carotid bifurcates after abdominal anesthesia. The common carotid artery was dissected free of the surrounding connective tissue and Ad-LDHA-WT, Ad-LDHA-K5R or Ad-LDHA-K76R (1×10^10^ pfu/mL) was suspended in 20 μL pluronic F127 gel (Sigma-Aldrich; 25% wt/vol) and applied around the carotid artery. Carotid arteries were harvested 14 days after ligation. Frozen arterial segments were sectioned at 5 μm and stained with hematoxylin and eosin (HE) and was examined by a light microscope (Nikon).

### Cell adenovirus infection

The VSMCs were infected with the above adenovirus (5×10^9^ pfu/mL) for 12 h, washed and incubated in serum-free medium without adenovirus for 12 h, then stimulated by PDGF-BB or crotonate, further cultured for 24 h.

### Western blot analysis

Cell samples were lysed in lysis buffer (50 mM Tris-HCl, pH 7.5, 150 mM NaCl, 1% NP-40, 1 mM EDTA, 0.11% sodium butyrate). Equal amounts of protein(20-100 μg) were separated on 10% SDS-PAGE gel and then transferred to PVDF membranes. After blocking with 5% nonfat milk for 2 hours at 37℃, membranes were incubated with specific primary antibody overnight at 4℃. Antibodies used in this study were obtained from the following sources: anti-LDHA antibody (Abclonal, A1146), anti-HA antibody (MBL, M180-3), anti-PCNA antibody (Proteintech, 10205-2-AP), anti-Kcr antibody (PTM Biolabs, PTM502), anti-Ub antibody (Santa Cruz Biotechnology, sc9133), anti-TOMM40 antibody (Abcam, ab272921), anti-tublin antibody (Proteintech, 10094-1-AP), anti-β-actin antibody (Cell Signaling Technology, #4970). Membranes were fourth washed with Tris-buffered saline containing 0.1% Tween 20 (TBST) and incubated with a HRP-conjugated secondary antibody (1:10000) for 2 hours. Immunoblots were visualized with the ECL detection system and quantified with ImageStudio software.

### siRNA transfection

The cultured VSMCs were grown to 50%-60% confluence, and then transfected with specific duplex siRNA, siLDHA, 5’-AGCAAAGAUUAUAGUGUGA-3’ and 5’-UCA CACUAUAAUCUUUGCU-3’, sicon, 5’-UUCUCCGAACGUGUCACGUTT-3’ and 5’-ACGUGACACGUUCGGAGAATT-3’ using Lipofectamine RNAiMIX reagent (Invitrogen) according to the manufacturer’s protocol. At 6 hours after transfection, VSMCs were incubated in a medium containing 5% FBS for 18h, then stimulated by PDGF-BB for 24 h.

### Immunofluorescence staining

VSMCs were fixed with 4% paraformaldehyde solution for 30 min at room temperature, washed twice with PBS for 5 min, and permeabilized with 0.1% Triton X-100 for 15 min. After blocking for 1.5 hours in 10% normal goat serum in a humidified chamber at 37℃, the cells were incubated for 3 hours at room temperature with rabbit anti-LDHA (1:50). Cells were washed 4 times with PBS and further stained with FITC-conjugated goat anti-rabbit secondary antibodies (1:100) for 2 hours at 37℃, followed by counterstaining with DAPI. For tissue immunofluorescence staining, the fixed carotid artery sections were washed three times for 5 min in the PBS and blocked in 5% goat serum for 2 hours at 37℃. Samples were incubated with anti-LDHA (1:100) and anti-PCNA (1:100) overnight at 4℃, followed by stained with secondary antibody conjugated with different fluorophore for 1 h at room temperature. Images were then observed on the confocal microscopy.

### Immunoprecipitation assay

The harvested cells were lysed as described above. Approximately 600 μg of the clarified cell lysate was incubated with 5μL indicated antibodies for 2 hours with rocking at 4℃, followed by incubation with 50μL protein A/G agarose overnight at 4℃. After 24 hours incubation, immunoprecipitates were collected by centrifugation and washed three times with 800 µL NP-40 buffer rotating for 20 minutes each time at 4℃. The immunoprecipitated protein then was supplemented with SDS-PAGE loading buffer, followed by Western blot analysis.

### Lactate production assay

Lactate levels were measured using a lactate colorimetric assay kit (BioVision) according to the manufacturer’s protocol. The absorbance at 450 nm was recorded using a BioTek microplate reader and normalized to protein concentration.

### Cell viability assay

Cells were seeded into 96-well plates and added CCK-8 test fluid after different treatment. Then cell viability were determined using CCK-8 kit (Abcam). After incubating for 1 h at 37℃, the absorbance was recorded at a wavelength of 450 nm in a microplate reader.

### Cell counting

The cells were digested with trypsin following diverse treatment, collected into 1.5mL EP tubes and washed with PBS, and then re-suspended with appropriate PBS. 10 µL cell suspension was added to the cell counting plate for determination.

### LDHA glutaraldehyde crosslink

Cells were lysed as described above. After protein quantification, equal amount of protein solution was added to a tube and replenished to 500 µL with NP40 lysate, 50% glutaraldehyde stock solution was added to the mixture, the final concentration of glutaraldehyde was 0.05%, and fixed on ice at 4℃ for 10 min. Then the fixation was terminated with 5× Launmi buffer and boiled for 5 min. Western blot analysis was performed as previously introduced.

### Mitochondrial extraction

VSMCs were serum-starved for 24 hours following adenovirus infection and PDGF-BB stimulation. Mitochondrial fractions were acquired using Mitochondrial Extraction kit (Invent Biotechnology) referring manufacturers’ recommendations.

### Mitochondrial staining

VSMCs were transferred to the circular plates, when the cells grew to appropriate density, the medium was replaced with DMEM containing Mito-Tracker (Invitrogen) and was preheated at 37 ℃ in advance. The live cells were stained for 45 min, then fixed with 4% paraformaldehyde at 37℃ for 30 min. Immunofluorescence staining was performed as above.

### Wound-healing assay

VSMCs were plated in six-well plates at appropriate density, after cell adherence and different treatment, unattached cells were rinsed with medium, a artificial wound was produced through scratching VSMCs with a 200ul sterilized pipette tip. Cell migration was then monitored by microscopy.

### Ni-NTA purification

Cells was lysed in Ni-agarose lysis buffer (50 mM NaH_2_PO_4_, 300 mM NaCl, 5mM imidazole, 0.05% Tween 20). Protein was purified by Ni-NTA purification kit (Invitrogen) according to manufacturers’ protocols and western blot was performed.

### Statistical analysis

All data analysis were performed by SPSS version 16.0. The data were presented as the means ± SEM. Statistical comparison of paired data was carried out using Student’s *t-*tests. The difference among groups were assessed by one-way or two-way analysis of variance (ANOVA). *P* <0.05 was considered statistically significant.

### Data availability

This study includes no data deposited in external repositories.

## Acknowledgements

We thank the members of our laboratories for comments and suggestions on this work.

## Author contributions

LH-D designed and directed the project; ZH-C performed the research with assistance from SH-C, ZY-R, HM-J and ZK-H; LH-D and ZH-C wrote the paper.

